# The emergence of Fluoroquinolone resistant and Extended-spectrum beta-lactamase-producing *Salmonella spp.* isolated from Poultry in Pakistan

**DOI:** 10.1101/2024.10.13.618098

**Authors:** Rimsha Irfan, Bushra Uzair, Eid Nawaz Khan, Abdullah Saeed

**Affiliations:** Department of Biological Sciences, International Islamic University, Islamabad 44000, Pakistan; Eid Nawaz Khan, senior Lab technologist at the National Veterinary Laboratory (NVL), Islamabad, Pakistan; Department of Internal Medicine, University of Michigan, Ann Arbor, MI 48109, United States

**Keywords:** Fluoroquinolone, Beta-lactamase, *Salmonella*, Poultry, Antimicrobial-resistance

## Abstract

The rise of antimicrobial-resistant and Extended Spectrum Beta Lactamases producing *Salmonella* strains in poultry is a severe health threat worldwide, particularly in developing countries like Pakistan. The current study aimed to investigate the isolation and identification of Cefotaxime and Ciprofloxacin-Resistant *Salmonella* strains originating from poultry. During this study, 78 (26.0%) *Salmonella* strains were isolated from 300 diverse poultry samples. The isolated 78 *Salmonella* strains were identified as *S. pullorum*, *S. gallinarum*, *S. enteritidis*, *S. enterica*, *S. paratyphi*, *S. typhimurium*, *S. typhi*, *S. arizonae,* and 19% other *Salmonella spp*. by API 20E method. The antibiotic susceptibility profile showed that 61% of MDR *Salmonella* strains were isolated from diverse poultry samples. MDR rates were high in serovars *S. enterica*, *S. typhimurium*, *S. enteritidis*, *S. arizonae,* and *S. gallinarum*. Co-resistant to cefotaxime and ciprofloxacin in 56.4% of *Salmonella* strains was observed. Among 44 phenotypically ESBL-positive *Salmonella* strains, 36 (81.8%) carried *bla* CTX-M and 38 (86%) carried *bla* CTX-M-1. A High incidence rate of *bla* CTX-M and *bla*-CTX-M-1 was observed among *S. enteritidis* (19% and 21%), *S. enterica* (11% and 16%), *S. typhimurium* (13% and16%) and *S. typhi* (3% and 5%). Among 29 phenotypically MBL-positive *Salmonella* strains, 13 (44.8%) carried *bla* VIM and 6 (20.6%) carried *bla* IMP. The 67% prevalence of *gyr* A was observed in fluoroquinolone-resistant *Salmonella* strains. The most prevalent fluoroquinolone strains were *S. enteritidis* (19%), *S. typhimurium* (19%), *S. typhi* (13%), and *S. gallinarum* (12%). Fluoroquinolone-resistant, MBL, and ESBL-producing *Salmonella strains* from poultry are a matter of great concern for both livestock and public health, demonstrating the dissemination risk of these microorganisms through the food chain.

## 1. Introduction

*Salmonella spp*. is one of the leading agents responsible for food-borne infections and diseases in humans, commonly in old-age and immune-compromised patients. Contaminated or uncooked food and poultry-origin products are the most common infection sources (Asif et al., 2017; Lamichhane et al., 2024).

Antimicrobial resistance is a growing problem in both human and veterinary treatment. The advent of antibiotic-resistant *Salmonella* strains is more virulent as they have raised public concern. The rise of antimicrobial resistance in *Salmonella* towards antibiotics such as ampicillin, cotrimoxazole, and chloramphenicol, can further complicate the cure and administration of enteric fever treatments (Hamdulay et al., 2024). Resistant Strains that have been detected in some clinical first-line antimicrobial drugs applied in the cure of severe *Salmonella* infections (Lamichhane et al., 2024).

ESBLs and MBLs producing Enterobacteriaceae pose a risk to human health (Bhandari et al., 2024). As in the cure of infections caused by Enterobacteriaceae in people, the β-lactam antimicrobial agents, especially cephalosporin and carbapenems, are the drugs of choice (Bologna et al., 2024). Treatment with antimicrobial drugs is life-saving, but the current resistance against traditional antibiotics has been directed to inadequate medication options (Singh et al., 2024).

Resistance to Cephalosporins is common in Enterobacteriaceae because of the production of broad-spectrum beta-lactamases, such as extended-spectrum-beta-lactamases (ESBLs) and Metallo-beta-lactamases (MBLs). Metallo-beta-lactamases (MBLs) are broad-spectrum zinc enzymes that are capable of in-activating most β-lactam antibiotics of clinical use, mainly Carbapenems which are known to be the most operative anti-biotics against *Salmonella spp*. Resistance to Carbapenems arises due to a decrease in anti-biotics absorption that occurs by lack an outer-membrane porin, reduction in permeability of outer-membrane, and MBL- production (Ghazaei, 2019; Zakhour et al., 2024). Fluoroquinolone resistance mostly occurs due to point mutations in those regions of gyrase genes (*gyr* A and *gyr* B), which determine quinolone resistance. Over the decade, broad-spectrum beta-lactam enzymes have been commonly confirmed in the microbiota of food animals (Akhlaghi et al., 2024). In Enterobacteriaceae including *Salmonella* species, CTX-M (Cefotaximase) kind ESBLs occur as highly prevalent beta-lactamase and cause human health problems (Tetteh et al., 2024). Resistance to ESBLs in bacteria is mainly based on plasmid-mediated production of enzymes that inactivate the antibiotics by hydrolyzing their β-lactam ring (Nasrollahian et al., 2024).

Bacteria can gain resistance to antimicrobial agents primarily by acquisition of mobile genetic elements and mutation in chromosomes such as plasmids by horizontal gene transmission (Vos et al., 2024). Antibiotic-resistant bacteria can be legitimately transferred through the food chain or transmit their antimicrobial resistance to human pathogens by portable genetic elements (Solanki and Das, 2024). To control this risk, it is necessary to monitor antimicrobial resistance in *Salmonella* strains (Ajayi et al., 2024). In Pakistan, no regulation or management for the suppression of *Salmonella* prevalence in poultry, and accurate diagnosis and targeted antibiotic treatment are still lacking (Harun et al., 2024; Wajid, Awan. The study aimed to determine the prevalence of *Salmonella* in poultry chains and to update knowledge on the prevalence of the ESBL and MBL-producing *Salmonella spp.* diversity in the poultry chain and their molecular characterization.

## 2. Materials and Methods

This study was carried out in Islamabad, the capital territory of Pakistan, at 60 poultry shops and farms in and around three different cities (Wah Cantt, Rawalpindi, and Islamabad). A total of 300 diverse poultry samples (100 from Chicken meat, 100 from Chicken ceca, and 100 from the Poultry environment) were collected and transported to the laboratory following the protocols.

### 2.1. Bacterial strains isolation and identification

The collected poultry samples were processed for isolation of *Salmonella* strains. To isolate bacteria, 15mg of each collected sample was added in Buffered peptone water at 1:10 for the pre-enrichment and incubated at 37°C for 24 hours. Afterward, *Salmonella’s* selective isolation was done by streaking a loopful culture on *Salmonella Shigella* (SS) agar and incubating them at 37°C for 24 hours.

The species-level identification was done by several biochemical methods (Gram Staining, Indole, Methyl-Red, Voges-Proskauer, citrate, and urease) and further confirmed by the API 20E system (BioMerieux^TM^ USA). H_2_S gas production and motility tests in the Sulfide-indole-motility medium were examined. In total 39 *Salmonella* strains were obtained from diverse poultry samples. Out of 39 *Salmonella* isolates 8 different serotypes were identified.

### 2.2. Screening for ESBLs and MBLs

*Salmonella* suspected colonies were inoculated on XLD agar plates comprising cefotaxime (4 μg/mL) and commercial HiCrome™ ESBL Agar Base (HiMedia, India) for screening the ESBL-producing *Salmonella spp*. For the screening of metallo-beta-lactamase-resistant *Salmonella*, the XLD agar supplemented with imipenem (1 μg/mL) was used. The plates were aerobically incubated at 37°C for 24 hours. All isolates, including the ESBL and MBL-producing *Salmonella spp.,* were preserved in brain heart infusion (BHI) broth and glycerol in cryovials and stored at −20°C.

### 2.3. Antibiotic Susceptibility Testing

Antimicrobial susceptibility for all the identified strains was determined by using the agar disc diffusion method following the CLSI recommendations. A total of 15 antimicrobial agents were tested: amoxicillin-clavulanic acid (AUG), cefotaxime (CTX), ceftazidime (CAZ), imipenem (IMP), meropenem (MEM), ceftriaxone (CRO), cefepime (FEP), gentamicin (CN), trimethoprim-sulfamethoxazole (SXT), ciprofloxacin (CIP), tetracycline (TE), azithromycin (AZM), aztreonam (ATM), cefoxitin (FOX), Piperacillin-Tazobactam (TZP). The results of the test strains were interpreted according to the CLSI guidelines.

Minimum inhibitory concentrations (MICs) of ciprofloxacin, cefotaxime, and ceftriaxone for all Salmonella isolates were determined by the Epsilometer strip (E-strip) test method. The antimicrobial agent concentration, at which the edge of the inhibition ellipse intersects the side of the E-strip, was taken as the MIC value. *Escherichia coli* ATCC 25922 was used as a control for MIC testing. MIC breakpoint results of the test strains were interpreted with the CLSI guidelines.

### 2.4. Phenotypic detection for ESBLs and MBLs

The ESBL production was detected through disc diffusion synergy test (DDST) using discs containing ceftazidime (CAZ) and cefotaxime (CTX) alone and with clavulanic acid in proximity to amoxicillin-clavulanic acid (AUG). A zone of inhibition was noted. The differences between inhibition zone diameters with single disc and combination discs were measured according to CLSI guidelines, and any isolate showing a difference of >5mm was considered as ESBLs producing bacteria (Wajid, Awan, et al., 2019).

MBL production was detected through a combined disc test (CDT) using discs containing meropenem (MEM) alone and meropenem discs impregnated with 0.5μg EDTA. Zones of inhibition were noted. A difference of ≥7mm between the inhibition zones of the meropenem discs tested alone compared to disks with EDTA infers MBL production phenotypically (Ghazaei, 2019).

### 2.5. Genomic DNA Extraction and Genotyping

The genomic DNA of bacteria was extracted by boiling method (Kim et al., 2024). Isolates with an ESBL and MBL phenotype were further examined by PCR to detect the presence of *bla* CTX-M-1 *bla* VIM and *bla* IMP. The fluoroquinolone-resistance-related *gyr*A gene was amplified by PCR in all ciprofloxacin-resistant *Salmonella* isolates. All primers and conditions for PCR amplification are provided in Table 1. The amplified PCR products were visualized in UV light after gel electrophoresis using 1.5 % agarose gel to obtain a better resolution of the amplified product, whose size was checked with standard molecular weight markers.

**Table 1.**
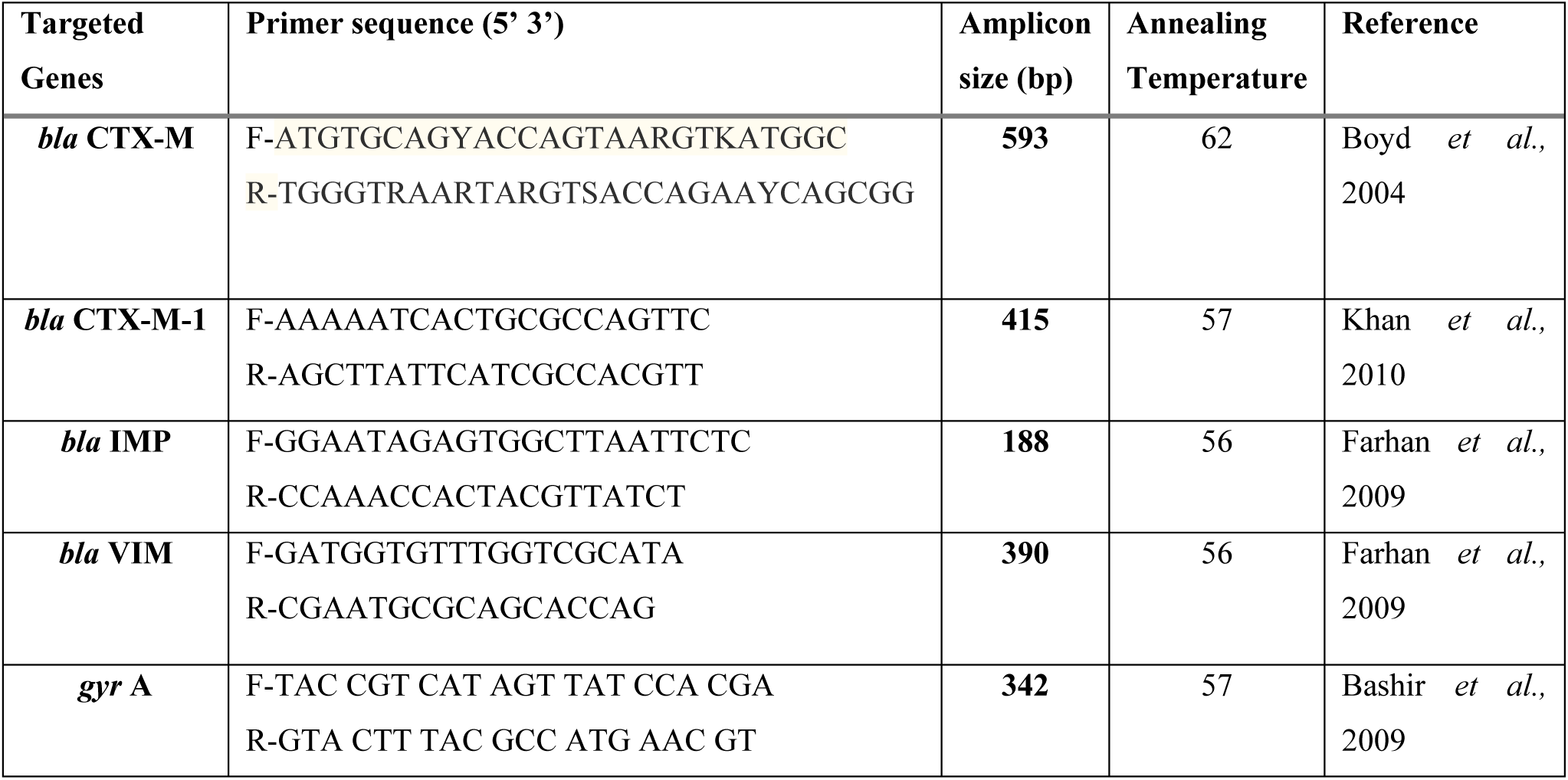
Primers used for amplification of ESBL, MBLs, and Fluoroquinolone resistance-related genes.

## 3. Results

Our study detected 78 (26.0%) CTX and CIP-resistant *Salmonella* isolates from 300 diverse poultry samples. API 20 E system, along with biochemical confirmation, gives species profiles of 62 *Salmonella* strains, characterizing them as *S. typhimurium* (*n*=6), *S. typhi* (*n*=4), *S. arizonae* (*n*=10), *S. pullorum* (*n*=10), *S. enteritidis* (*n*=8), *S. gallinarum* (*n*=10), *S. enterica* (*n*=6), *S. paratyphi* (*n*=8). At the same time, 16 isolates were confirmed as *Salmonella* strains (99.3 %).

Antimicrobial susceptibility tests showed a significant amount of antibiotic resistance in *Salmonella* strains. Out of 78 isolates, 44(56.4%) *Salmonella* strains were co-resistant to cefotaxime and ciprofloxacin. Resistance to 5-9 antibiotics was detected in 61.5% (48/78) isolates. Notably, MDR rates were high in serovars *S. enterica*, *S. typhimurium*, *S. enteritidis*, *S. arizonae,* and *S. gallinarum* isolated from diverse poultry samples. More of the serovars isolated from chicken ceca showed MDR 45.8% (*n*=22/48), serovars from chicken meat showed MDR 29.1% (*n*=14/48) than did the serovars isolated from poultry environment showed MDR 25% (*n* =12/48).

The phenotypic profile for ESBL production, out of 78 *Salmonella* isolates, 44 (56.4%) of the isolates were positive for ESBL production. Regarding ESBL-producing *Salmonella* isolates origin, 45.4% (20/44) were from the chicken ceca, 34.0% (15/44) from chicken meat, and 20.4% (9/44) from the poultry environment. The highest ESBL production rate was observed among the serovars *S. enterica*, *S. typhimurium,* and *S. enteritidis*. The phenotypic profile for MBL production, out of 78 *Salmonella* isolates, 29 (37.1%) of the isolates were positive for MBL production. Regarding MBL-producing *Salmonella* isolates origin, 44.8% (13/29) were from the chicken ceca, 31.0% (9/29) from chicken meat, and 24.1% (7/29) from the poultry environment.

The *Salmonella* isolates that showed phenotypic resistance characteristics against Cefotaxime, and Ciprofloxacin was further assessed to confirm their genotypic resistance. The isolates showed positivity for the genes studied. In the current study, an 81.8% prevalence of *bla* CTX-M was observed in *Salmonella* strains, while 86% *bla* CTX-M-1 positive *Salmonella* strains were observed in current research.

The current study detected 44.8% (13/29) bla VIM and 27.5% (8/29) bla IMP in phenotypically MBL-positive *Salmonella* strains. The fluoroquinolone resistance determinants were analyzed among 48-CIP-resistant *Salmonella* strains in this study. The prevalence of *gyr* A, 67%, was observed in fluoroquinolone-resistant *Salmonella* strains.

**Figure 2.**
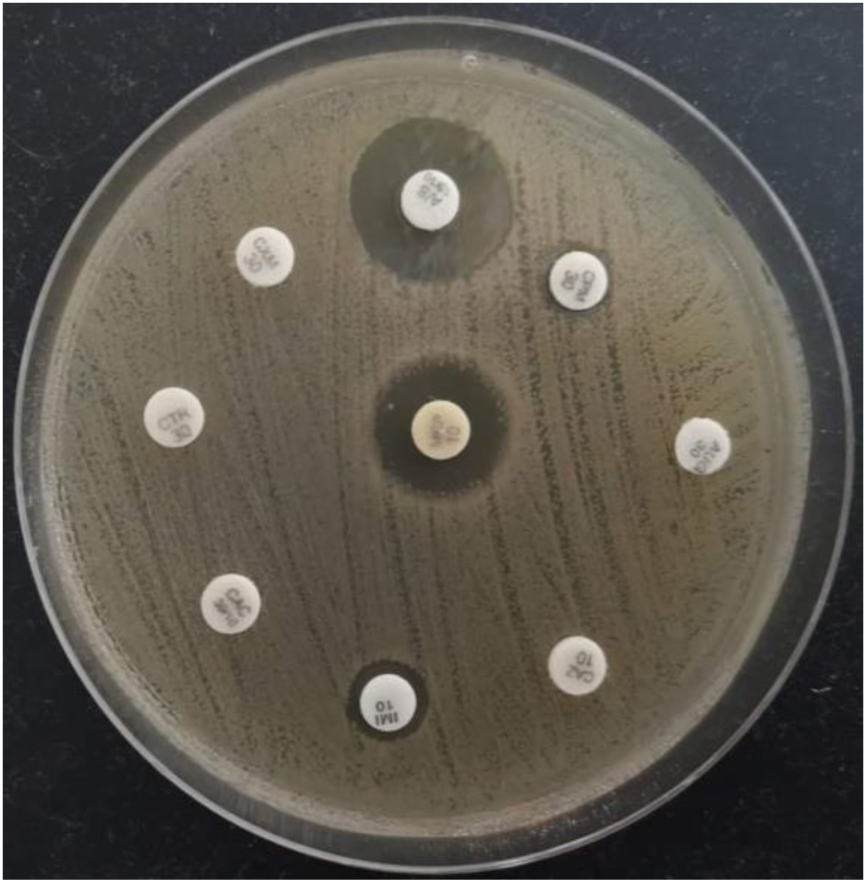
Antibiotic sensitivity patterns of MDR *Salmonella enterica*

**Figure 2.**
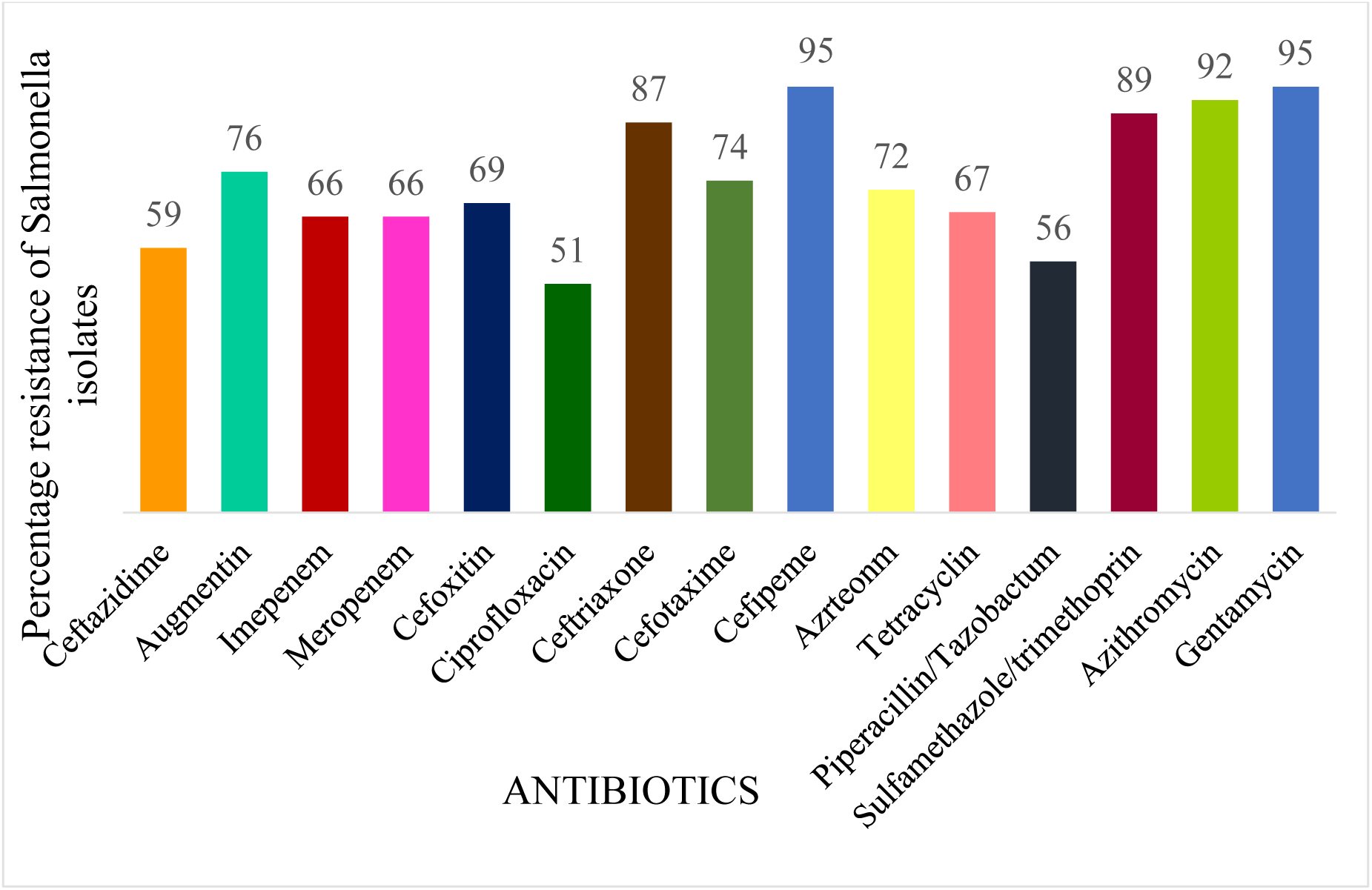
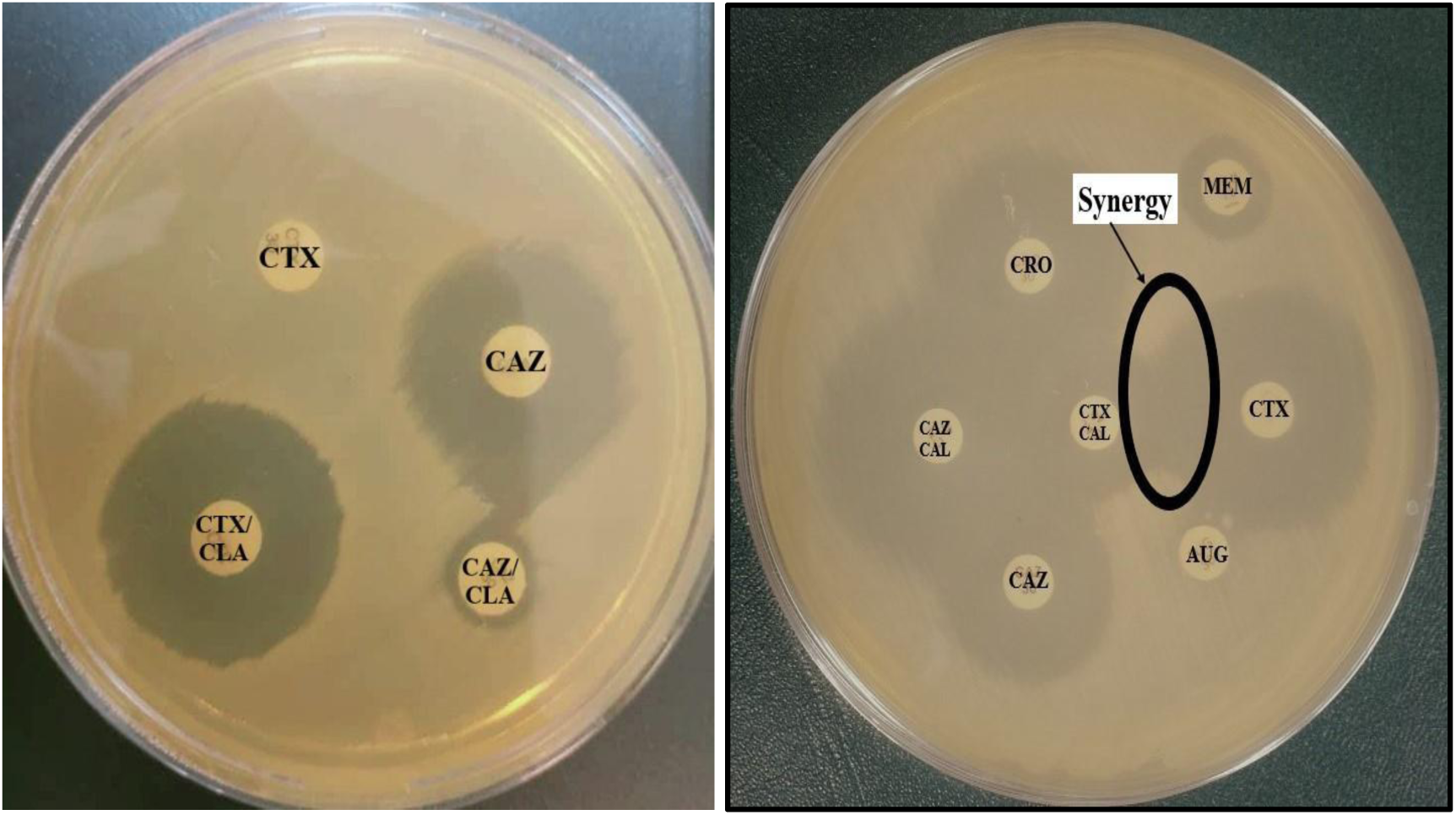
Phenotypic detection of ESBL-producing *Salmonella* by Disc Diffusion Synergy Test

**Figure 3.**
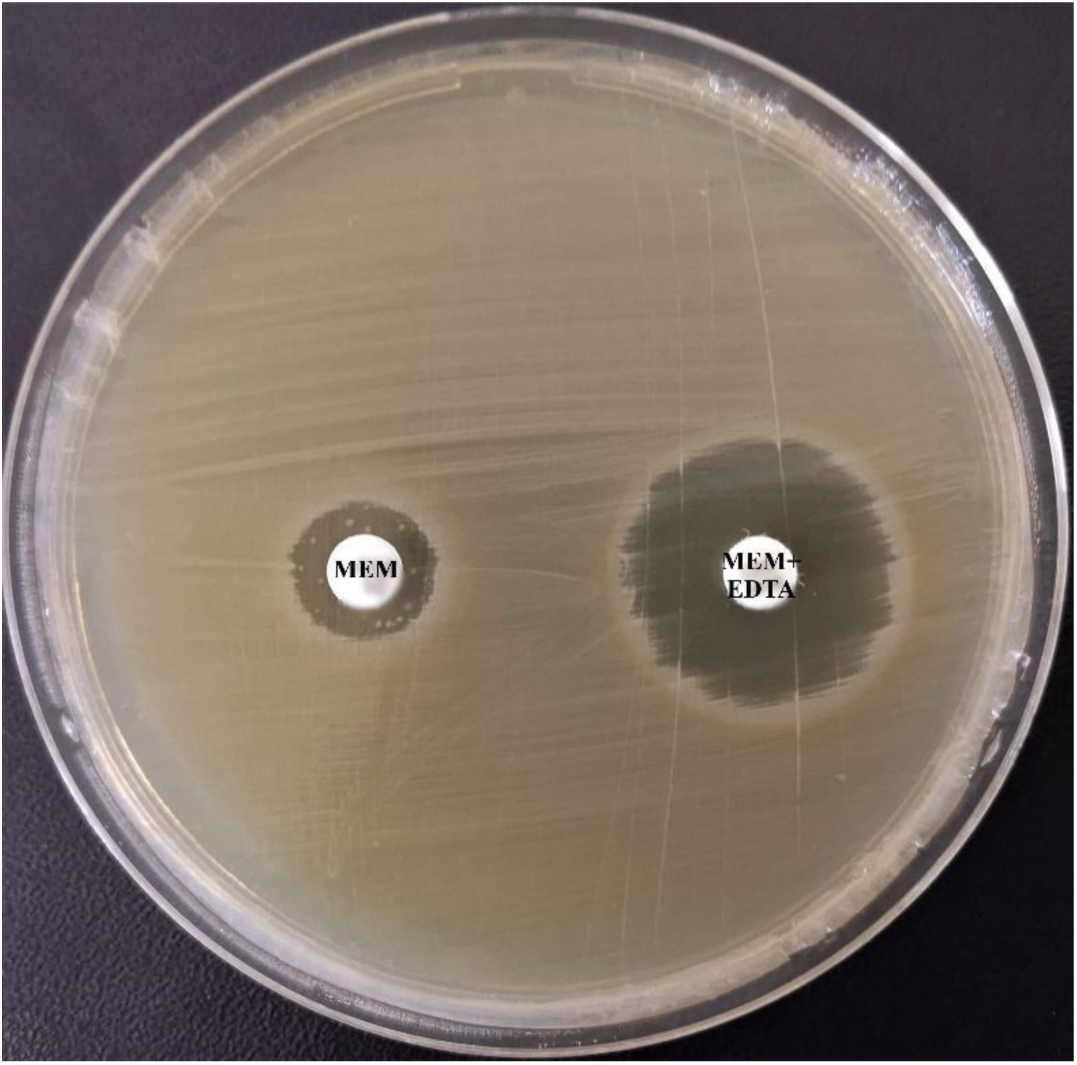
Phenotypic detection of MBL-producing *Salmonella* by Combined Disc Synergy Test

**Figure 3.**
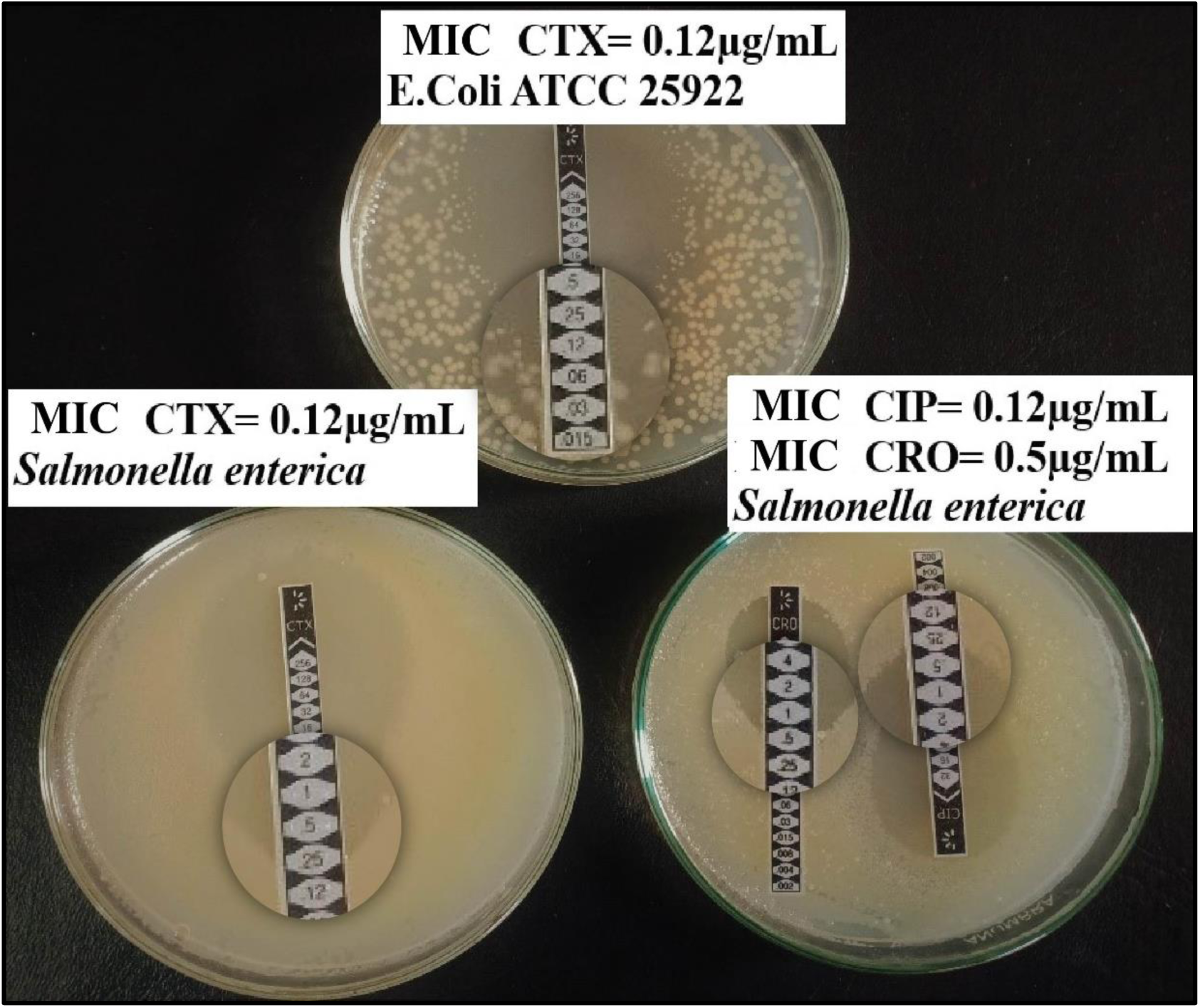
MIC Determination of *Salmonella enterica* against Cefotaxime (CTX), Ciprofloxacin (CIP), and Ceftriaxone (CRO)by E-Strip Test.

### 3.1. PCR Analysis of *bla* CTX-M-1, *bla* VIM, *bla* IMP, and *gyr* A genes

**Table 3.**
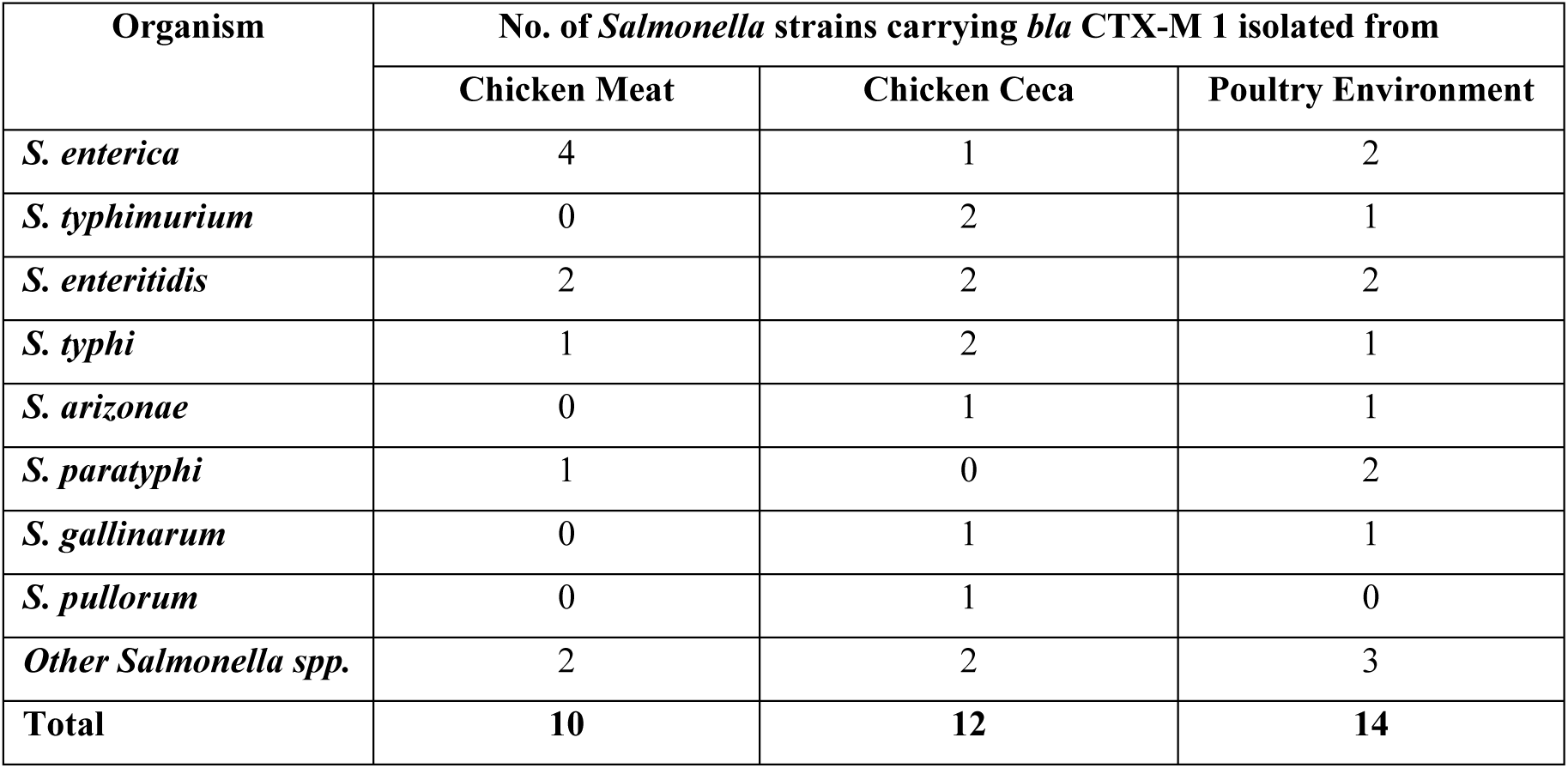
Prevalence of *bla* CTX-M (ESBLs encoding gene) in *Salmonella spp.* isolated from diverse poultry samples.

**Table 4.** Prevalence of *bla* CTX-M-1 (ESBLs encoding gene) in *Salmonella spp.* isolated from diverse poultry samples.

| Organism | No. of *Salmonella* strains carrying *bla* CTX-M 1 isolated from |  |  |
| --- | --- | --- | --- |
|  | Chicken Meat | Chicken Ceca | Poultry Environment |
| *S. enterica* | 6 | 3 | 0 |
| *S. typhimurium* | 0 | 2 | 3 |
| *S. enteritidis* | 4 | 5 | 2 |
| *S. typhi* | 1 | 2 | 1 |
| *S. arizonae* | 1 | 1 | 0 |
| *S. paratyphi* | 0 | 2 | 0 |
| *S. gallinarum* | 0 | 0 | 0 |
| *S. pullorum* | 0 | 1 | 0 |
| *Other Salmonella* spp. | 1 | 0 | 3 |
| **Total** | **13** | **16** | **9** |

**Table 5.** Prevalence of *gyr* A (Fluoroquinolone-resistant related gene) in *Salmonella* strains isolated from diverse poultry samples.

| Organism | No. of *Salmonella* strains carrying *gyr* A isolated from |  |  |
| --- | --- | --- | --- |
|  | Chicken Meat | Chicken Ceca | Poultry Environment |
| *S. enterica* | 2 | 0 | 0 |
| *S. typhimurium* | 0 | 2 | 1 |
| *S. enteritidis* | 0 | 2 | 1 |
| *S. typhi* | 1 | 1 | 0 |
| *S. arizonae* | 1 | 0 | 1 |
| *S. paratyphi* | 0 | 0 | 0 |
| *S. gallinarum* | 0 | 2 | 0 |
| *S. pullorum* | 0 | 0 | 0 |
| *Other Salmonella* spp. | 0 | 0 | 2 |
| **Total** | **4** | **7** | **5** |

**Table 6.** Prevalence of *bla* IMP in *Salmonella* strains isolated from diverse poultry samples.

| Organism | No. of *Salmonella* strains carrying *bla* IMP isolated from |  |  |
| --- | --- | --- | --- |
|  | Chicken Meat | Chicken Ceca | Poultry Environment |
| *S. enterica* | 1 | 0 | 0 |
| *S. typhimurium* | 0 | 0 | 1 |
| *S. enteritidis* | 0 | 0 | 1 |
| *S. typhi* | 1 | 1 | 0 |
| *S. arizonae* | 1 | 0 | 0 |
| *S. paratyphi* | 0 | 0 | 0 |
| *S. gallinarum* | 1 | 1 | 0 |
| *S. pullorum* | 0 | 0 | 0 |
| *Other Salmonella spp.* | 0 | 0 | 0 |
| **Total** | **4** | **2** | **2** |

**Table 7.** Prevalence of *bla* VIM in *Salmonella* strains isolated from diverse poultry samples.

| Organism | No. of *Salmonella* strains carrying *bla* VIM isolated from |  |  |
| --- | --- | --- | --- |
|  | Chicken Meat | Chicken Ceca | Poultry Environment |
| *S. enterica* | 0 | 1 | 1 |
| *S. typhimurium* | 0 | 0 | 1 |
| *S. enteritidis* | 0 | 0 | 1 |
| *S. typhi* | 1 | 1 | 0 |
| *S. arizonae* | 1 | 0 | 0 |
| *S. paratyphi* | 0 | 1 | 0 |
| *S. gallinarum* | 1 | 1 | 0 |
| *S. pullorum* | 0 | 0 | 0 |
| *Other Salmonella spp.* | 1 | 2 | 0 |
| **Total** | **4** | **6** | **3** |

**Figure 4.**
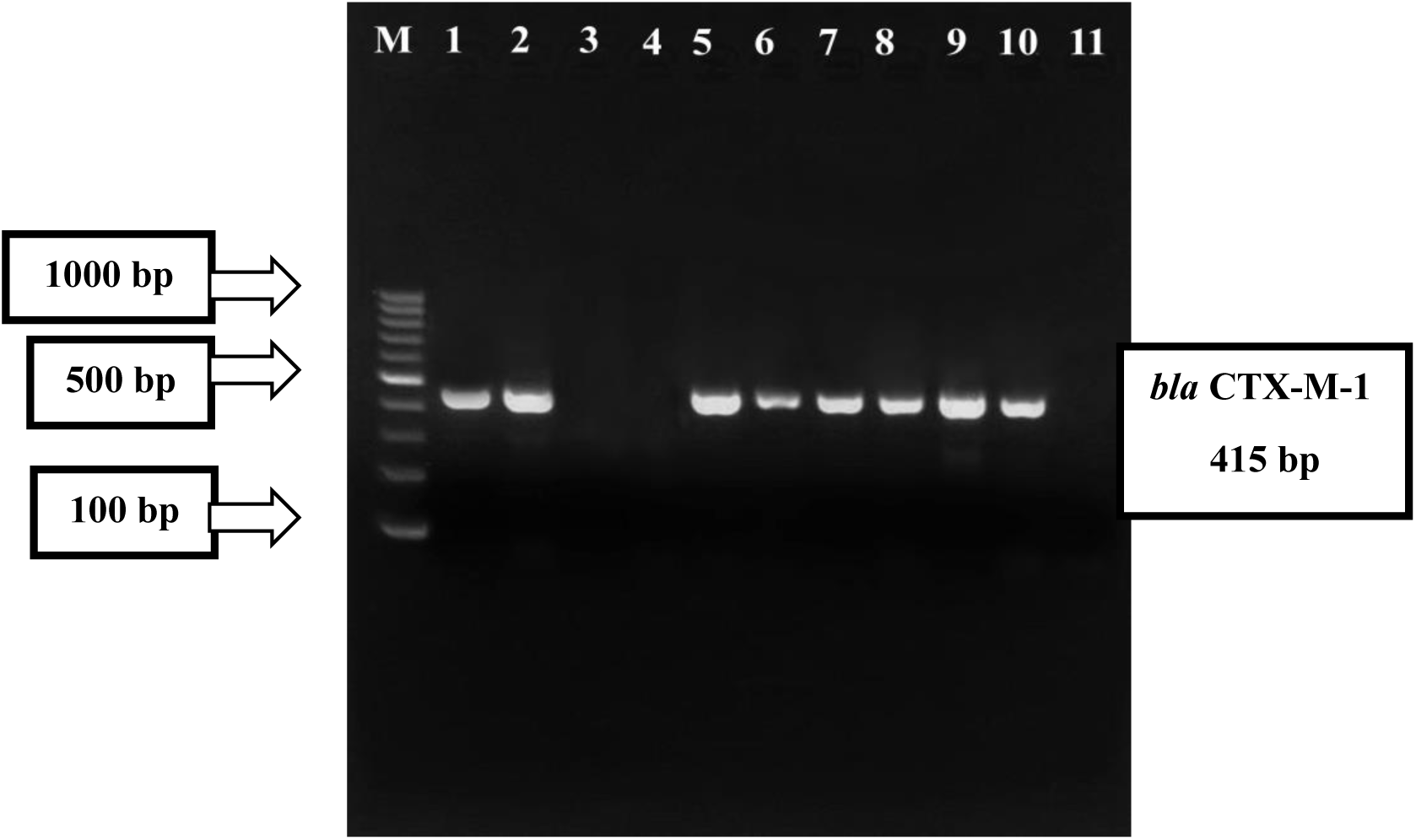
PCR-based detection of *bla* CTX-M-1 gene in phenotypically ESBL-positive *Salmonella spp.* **Key: Lane M:** 100bp Molecular Ladder **Lane 1,2,5-10:** *bla* CTX-M-1 Positive *Salmonella spp.* **Lane 3,4,11:** *bla* CTX-M-1 Negative *Salmonella spp*.

**Figure 5.**
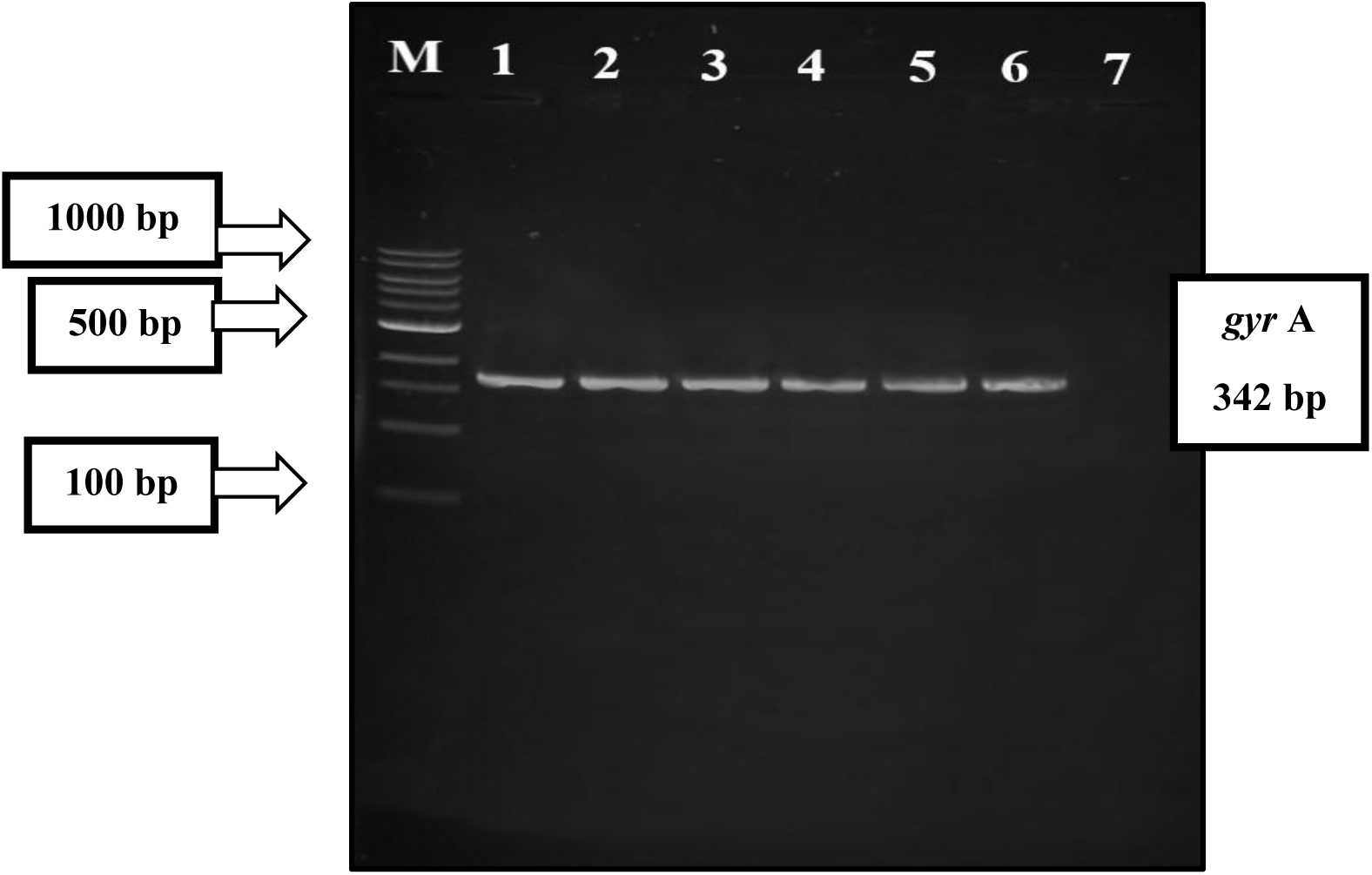
PCR-based detection of *gyr* A gene in phenotypically CIP-resistant *Salmonella spp.* **Key: Lane M:**100bp Molecular Ladder **Lane 1-6:** *gyr* A Positive *Salmonella spp.* **Lane 7:** *gyr* A Negative *Salmonella spp*.

**Figure 6.**
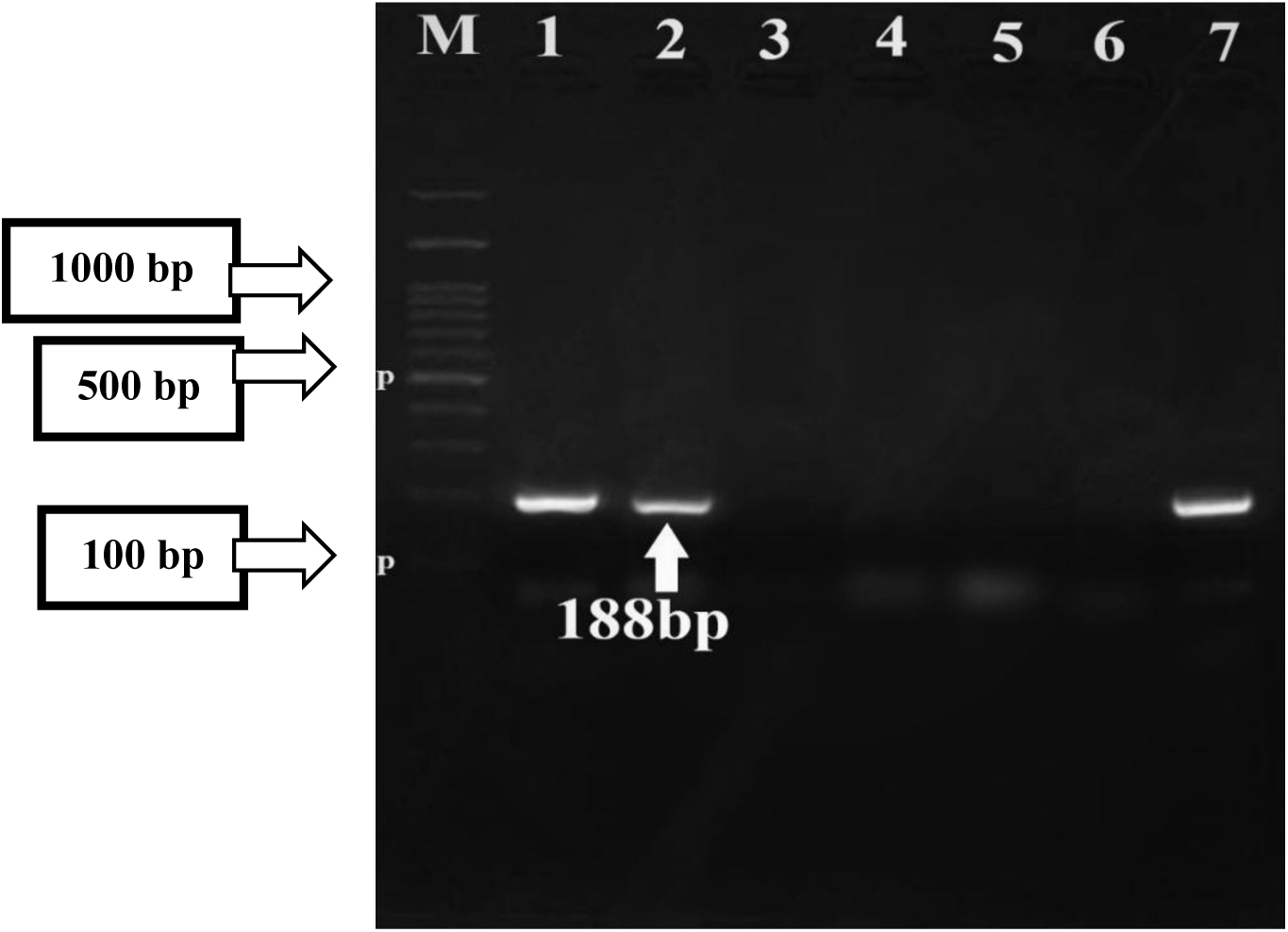
PCR-based detection of *bla* IMP gene in phenotypically MBL-producing *Salmonella spp.* **Key: Lane M:**100bp Molecular Ladder **Lane 1,2,7:** *bla* IMP Positive *Salmonella spp.* **Lane 3-6:** *bla* IMP Negative *Salmonella spp*.

**Figure 7.**
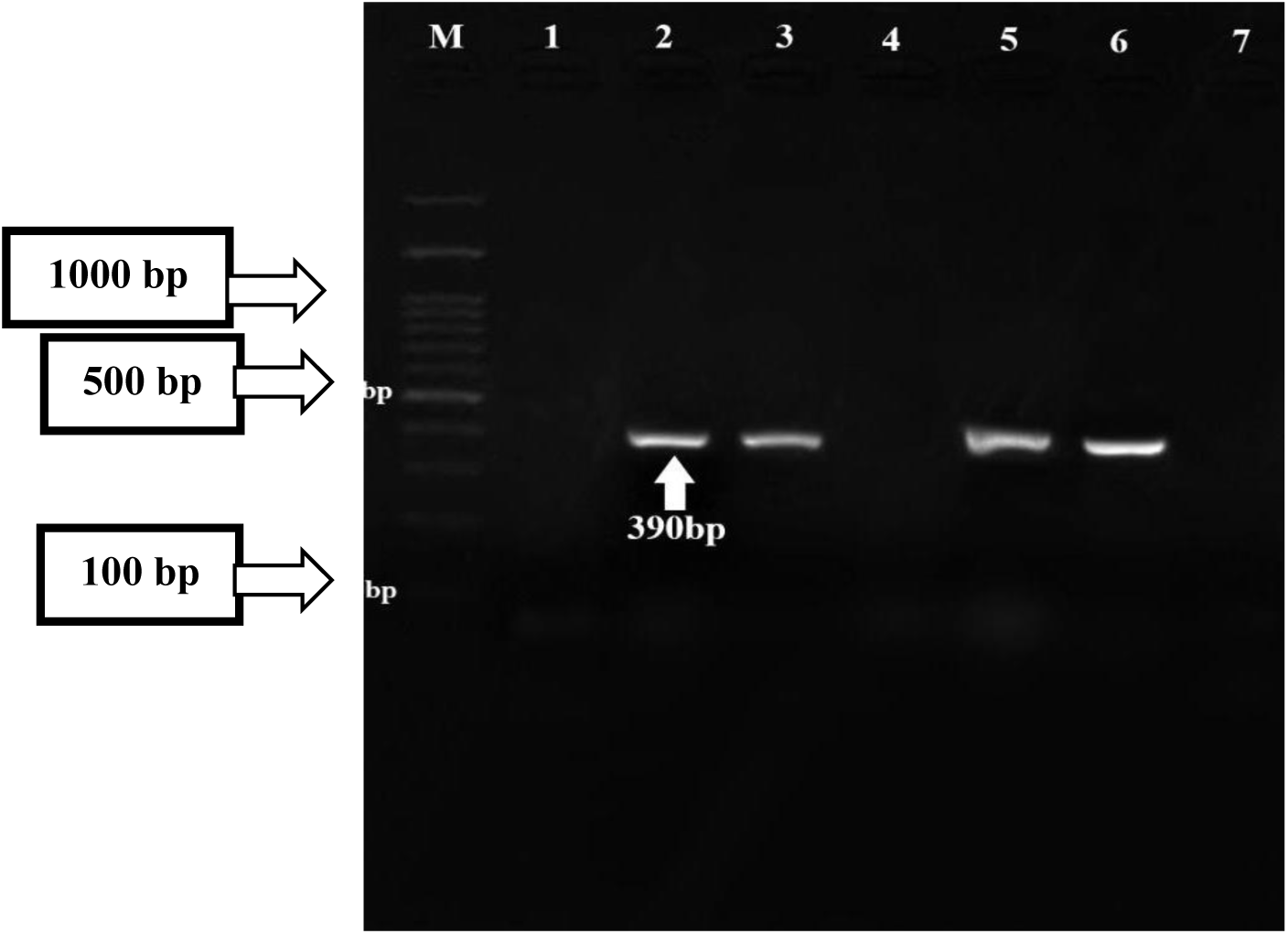
PCR-based detection of *bla* VIM gene in phenotypically MBL-producing *Salmonella spp.* **Key: Lane M:**100bp Molecular Ladder **Lane 2,3,5,6:** *bla* VIM Positive *Salmonella spp.* **Lane 1,4,7:** *bla* VIM Negative *Salmonella spp*.

## 4. Discussion

The increase in antibiotic-resistant *Salmonella* strains is more virulent as it has raised public concern, triggering a rise in the death rate of infected people. Studies have shown the incidence of MDR (Multi-Drug-Resistant) *Salmonella* in food and animal products. The rates of *Salmonella* infections have also increased. (Billah and Rahman, 2024; Wajid, Awan, et al., 2019; Wajid, Saleemi, et al., 2019).

To the best of our knowledge, this is the first time an attempt has been made to determine and isolate ESBL and MBL-producing and Fluroquinolone-resistant *Salmonella* strains from poultry in this region.

During this study, 300 samples of chicken meat, chicken ceca, and poultry environment from the cities of Islamabad, Rawalpindi, and Wah Cantt were collected. Among them, 78 (26.0%) were positive for *Salmonella* strains. From Wah Cantt, 22%, Rawalpindi, 26%, and Islamabad, 30% of *Salmonella* strains were isolated. The highest prevalence rate of *Salmonella* strains in poultry was observed in areas of Islamabad (30%).

The highest isolation rate of *Salmonella* strains was obtained from chicken ceca (44%) followed by chicken meat (25%). The isolated *Salmonella* strains were identified as *Salmonella pullorum* (14%), *Salmonella gallinarum* (13%), *Salmonella enteritidis* (10%), *Salmonella enterica* (8%), *Salmonella paratyphi* (10%), *Salmonella typhimurium* (8%), *Salmonella typhi* (5%), *Salmonella arizonae* (13%) and 19% other *Salmonella spp*. by API 20 E method.

In Brazil, Pribul and colleagues found 33 different *Salmonella* strains isolated from food and environmental samples, among which *Salmonella typhimurium* 48% was the prevalent strain supported by *Salmonella enteritidis* 19%. This study showed that several antimicrobial-resistant *Salmonella* strains were isolated from diverse poultry samples. The highest resistance rates were found against gentamycin 95%, cefepime 95% and azithromycin 92%, followed by sulfamethoxazole/ trimethoprim 90% and ceftriaxone 87% (Pribul et al., 2017).. This outcome is inconsistent with the data from another study in which all the tested *Salmonella* strains were resistant to ceftriaxone 95% and sulfamethoxazole/ trimethoprim 70% (Ghazaei, 2018).

Suresh *et al*. reported that *Salmonella* strains were resistant to gentamicin ceftriaxone 57.1% and sulfamethoxazole/trimethoprim 71.1%. Interestingly, 22 (56.4%) *Salmonella* strains were co-resistant to cefotaxime and ciprofloxacin. Co-resistance among cefotaxime and ciprofloxacin has been previously described in *Salmonella* isolates (Suresh et al., 2019). In China, Bai *et al.,* 2015 observed that 68.7% of *Salmonella* strains of chicken meat origin were co-resistant to cefotaxime and ciprofloxacin (Bai et al., 2015). Similarly, *Salmonella* from poultry origin has been found to be co-resistant to ciprofloxacin and cefotaxime, which was quite lower-rate resistance than the present study’s findings (Castro-Vargas et al., 2020).

Intermediate resistance to cefotaxime, meropenem, and aztreonam was shown by all isolates. Resistance to 5-9 antibiotics was detected in 61% of isolates. Current study findings are in line with a study performed in Kohat, Pakistan, in which 54.2% MDR *Salmonella* strains were obtained from poultry samples (Acharya et al., 2023). In another recent research in Faisalabad, Pakistan, 26% of MDR *Salmonella* isolates in poultry were reported (Wajid, Awan, et al., 2019). Notably, MDR rates were high in serovars *S. enterica*, *S. typhimurium*, *S. enteritidis*, *S. arizonae*, and *S. gallinarum* isolated from diverse poultry samples. The results showed that 45.8% of MDR Salmonella strains were isolated from chicken ceca, 29% from chicken meat, and 25% from the poultry environment.

Notably, the high rate of antimicrobial-resistant *Salmonella* strains in the present study is because of the primary introduction and subsequent common usage of these antibiotics in veterinary medication and in humans in Pakistan. Immense inoculation of herds of animals by some farmers has been increased as a contributing aspect in the increase in antimicrobial resistance, as well as the lack of regulations and monitoring in the management of antimicrobial drugs in veterinary establishments.

ESBL-positive *Salmonella* strains have been reported in farm animals and food. Poultry is the most important cause of ESBL-producing *Salmonella*. Cephalosporins and fluoroquinolones were used to treat *Salmonella* infections after the emergence of MDR *Salmonella.* This study showed that ESBL-positive isolates displayed greater antimicrobial resistance than non-ESBL producers.

The current results show that 56.4% of the strains presented the phenotypic profile for ESBL production. The detection percentage of ESBL-producing *Salmonella* isolates in the present study was significantly higher than those of ESBL-production rate at 10.8% and 8.8% in *Salmonella* strains isolated from poultry stated in China (Qiao et al., 2017) and Pakistan (Wajid, Awan, et al., 2019) respectively.

In Egypt, Elhariri *et al*. reported 66.6% ESBL-positive Salmonella strains among broiler chicken and poultry environments, which is in line with the present study’s findings (Elhariri et al., 2020). Notably, the highest ESBL-production rate was observed among the serovars *S. enterica*, *S. typhimurium,* and *S. enteritidis*. In the present study, all these strains showed ESBL production. These rates are comparatively high with the findings reported from Iran, where 25% of *S. typhimurium* and 75% of *S. enteritidis* ESBL-producing strains were isolated (Ghazaei, 2018).

A study conducted in Nepal Bahartpur by Bajracharya *et al*. reported 62.5% MBL-positive *Salmonella* spp. Isolates (Bajracharya et al., 2023). Those *Salmonella* isolates that showed phenotypic resistance characteristics against Cefotaxime, and Ciprofloxacin was further assessed to confirm the genotypic resistance in them. The isolates showed positivity for the genes studied. The current study observed an 81.8% prevalence of bla CTX-M in *Salmonella* strains. While 86% *bla* CTX-M-1 positive *Salmonella* strains were observed in this research. A study conducted in Algeria by Djeffal *et al*., 2017 reported 66.7% of *bla* CTX-M in ESBL-producing *Salmonella* from poultry (Djeffal et al., 2017). The present study results are much higher comparatively with findings of a study performed in Iran showing that 69% *bla* CTX-M positive ESBL-producing *Salmonella spp.* in broiler (Haeri and Ahmadi, 2019).

de Souza *et al*. stated 55% *bla* CTX-M in phenotypically ESBL-positive *Salmonella* strains (de Souza et al., 2019). A very low prevalence rate of *bla* CTX-M (23%) had been reported in Ceftiofur-resistant *Salmonella* isolated from poultry (Denagamage et al., 2019). The present study detected a considerable prevalence of *bla* CTX-M-1 in *S. enteritidis* (21%), *S. enterica* (16%), *S. typhimurium* (16%), and *S. typhi* (5%).

The fluoroquinolone resistance determinants were analyzed among 48-CIP-resistant *Salmonella* strains in this study. The prevalence of *gyr* A with 67% was observed in fluoroquinolone-resistant *Salmonella* strains. The present study results coincide well with the research in Korea by Kim et al., who had reported a 62.3% prevalence of fluoroquinolone-resistance-related gene (*gyr* A) in *Salmonella* strains isolated from chicken (Kim et al., 2019). In contrast, in a similar approach, A study reported 29% (12/41) mutated *gyr* A detection in *Salmonella* isolates (Al-Gallas et al., 2021). In China, Chen *et al*. reported no mutations in *gyr* A were observed in Ciprofloxacin and ceftriaxone-resistant *Salmonella* serovars from chicken meat (Chen et al., 2019).

In the current study, the most prevalent strains were *S. enteritidis* (19%), *S. typhimurium* (19%), *S. typhi* (13%) and *S. gallinarum* (12%). Likewise, the presence of 44.8% (13/29) *bla* VIM and 27.5% (8/29) *bla* IMP was detected in *Salmonella* strains. The results are in line with the research in Iran by Ghazaei, who had reported a 31.57% prevalence of *bla* IMP and 57.89% *bla*VIM gene in *Salmonella enterica* isolated from poultry meat (Ghazaei, 2019).

In conclusion, the *Salmonella* stains present phenotypic and genotypic characteristics for Fluoroquinolones MBL and ESBL production, demonstrating the dissemination risk of these microorganisms through the food chain.

## 5. Conclusion

Third-generation cephalosporins, used to treat enteric fever and other infections caused by *Salmonella* strains, may increase the incidence of CTX-M-producing *Salmonellae*. Decreased susceptibility to ciprofloxacin among *Salmonella* strains has also contributed to the increased use and subsequent development of resistance to cephalosporins among *Salmonellae*.

This study indicated a high ESBL, MBL, and fluoroquinolones burden in poultry. This signifies the role of the food of animal origin as a reservoir of MDR *Salmonella* that can affect human health. All major resistance determinants, including those that confer resistance against ESBLs, MBLs, and Fluoroquinolones, have been identified in *Salmonella* strains. In addition, the current study suggests the role of animal-based food as a source of MDR *Salmonella,* demanding veterinary supervision on antibiotic use for therapy purposes in poultry. The current data concerning the epidemiological spreading of MDR *Salmonella* strains between humans and poultry, this study provides valuable information correlated to the circulation of MRD- resistant *Salmonella* strains. This underscores the need to continue surveilling food-borne bacteria along the poultry chain. It has become progressively clear that antibiotic resistance will remain a significant barrier to challenge in the future. The discovery of gene modifications in beta-lactamase-producing microbes is important evidence for the proper and effective therapy of Salmonellosis.

## CRediT authorship contribution statement

**Rimsha Irfan:** Writing – review & editing, Writing – original draft, Methodology, Formal analysis, Data curation, Visualization, Validation, Conceptualization. **Bushra Uzair:** Writing – review & editing, Supervision, Resources, Project administration, Visualization, Conceptualization, Funding acquisition. **Eid Nawaz Khan:** Methodology, Investigation. **Abdullah Saeed:** Review & editing

## Acknowledgments

We are thankful to Dr. Muhammad Javed Arshad (senior scientific officer), Dr. Shumaila Manzoor (Laboratory Technologist), and Mr. Eid Nawaz Khan (Laboratory Technologist) for their guidance and support. The present study was approved by the Institution Review Board and Ethical Review Board of International Islamic University, Islamabad, Pakistan (Reg#330- FBAS/MSBT/F-17). This research received no external funding.

